# cellxgene: a performant, scalable exploration platform for high dimensional sparse matrices

**DOI:** 10.1101/2021.04.05.438318

**Authors:** Colin Megill, Bruce Martin, Charlotte Weaver, Sidney Bell, Lia Prins, Seve Badajoz, Brian McCandless, Angela Oliveira Pisco, Marcus Kinsella, Fiona Griffin, Justin Kiggins, Genevieve Haliburton, Arathi Mani, Matthew Weiden, Madison Dunitz, Maximilian Lombardo, Timmy Huang, Trent Smith, Signe Chambers, Jeremy Freeman, Jonah Cool, Ambrose Carr

## Abstract

Quickly and flexibly exploring high-dimensional datasets, such as scRNAseq data, is underserved but critical for hypothesis generation, dataset annotation, publication, sharing, and community reuse. cellxgene is a highly generalizable, web-based interface for exploring high dimensional datasets along categorical, continuous and spatial dimensions, as well as feature annotation. cellxgene is differentiated by its ability to performantly handle millions of observations, and bridges a critical gap by enabling computational and experimental biologists to iteratively ask questions of private and public datasets. In doing so, cellxgene increases the utility and reusability of datasets across the single-cell ecosystem.

The codebase can be accessed at https://github.com/chanzuckerberg/cellxgene. For questions and inquiries, please contact.

## Introduction

As biological datasets grow in size and complexity, complementary sets of expertise are required to drive scientific discovery. Computational skills and statistical awareness are required for managing, analyzing, and visualizing the data, while a rich knowledge of biological systems is crucial for asking insightful questions and interpreting results. Rich and complex biological datasets often require collaborative teams, which face the challenge of identifying tools to facilitate diverse researchers to offer their expertise and input.

Domain experts and bioinformaticians typically use different tools. Bioinformaticians often explore and visualize datasets through interactive computing environments, such as Jupyter and RStudio. These environments allow them to iterate through cross-sections and views of the data, which enables rapid exploration and provides context for observations. However, sharing this context and insight with experimental biologists and clinicians is highly inefficient because much of the information is lost in the translation to static, shareable outputs. Exchanges between collaborators are manual and frustrating, such as continuously emailing slightly-updated spreadsheets and flat figures back and forth.

This problem is consistent across many subfields, including single-cell biology. Datasets consist of tens of thousands of measurements (e.g., gene expression) across millions of individual observations (e.g., cells).

cellxgene (*cell-by-gene*) is a data exploration and visualization tool that enables rich, intuitive, collaborative interrogation of high-dimensional datasets at scale. The tool enables both bioinformaticians and experimentalists to load the full, interactive results of their computational analyses, which reduces communication burden by empowering experimental researchers to independently interrogate and annotate the data with minimal training. In doing so, cellxgene bridges the gap between technical and domain expertise, greatly facilitating the exploration and discovery process.

While many noteworthy visualization tools have been developed, cellxgene is differentiated by its adherence to best practices in information design and software engineering; a design tailored to biologists’ needs; and the ability to scale seamlessly to handle millions of observations and tens of thousands of measurements without sacrificing real-time interactivity. While the target domain of cellxgene is single-cell datasets, its internal architecture is domain-agnostic and generalizable to any matrix-form dataset. Within single-cell biology, early use has demonstrated the exploration of scRNA, snRNA, ATAC, single-cell spatial analysis, proteomics, and more.

## Core features and functionality

cellxgene focuses on making exploration and examination of analytical results simple and fast. The tool ingests (1) matrix-form datasets (e.g., expression matrices) consisting of observations (e.g., cells) as rows and features (e.g., gene expression or other abundance measurements) as columns; (2) continuous or categorical metadata in the same format; (3) a pre-computed, initial embedding of the cells in a low-dimensional space (e.g., PCA, tSNE, UMAP, or spatial coordinates). This format accommodates the output of most analysis toolchains in single-cell biology, such as Seurat, Scanpy, Bioconductor, or scVI. The results of any additional computational analyses, such as clustering, can be imported alongside other metadata.

The visualizations in cellxgene are intuitive and intentionally simple. The primary visualization is a scatterplot of cells embedded in two dimensions based on their gene expression values or spatial coordinates. This is complemented by histograms displaying the distribution of each feature and metadata field. Users can filter these visualizations by interacting with facets based on gene expression, metadata, and the embedding itself. The combinatorial nature of these affordances facilitates arbitrary exploration, cross-filtering, subsetting, coloring, and computationally re-embedding subpopulations of the data.

cellxgene enables researchers to identify relationships between subpopulations of cells, gene expression, and cell metadata. Based on these observations, users can then probe interesting facets of the data by subsetting, computing differential expression (e.g., t-tests), and re-embedding subpopulations of cells. Findings can be recorded and shared via exportable annotations.

## Common uses in single-cell biology

A single-cell biologist may begin by investigating whether the underlying structure of the data, represented by the position of observations in the embedding, is a result of technical variation. The researcher would accomplish this by coloring the scatterplot by donor, the replicate the cell was derived from, or another technical metadata field. This yields immediate insights into whether the data contains uncorrected technical variation (for instance, if donors were not evenly mixed in the embedding).

In addition, painting the scatterplot by a variable automatically renders its conditional distribution across each other variable (e.g., does each donor contribute cells to each cluster, or are some clusters donor-specific? is the expression for a tumor suppressor gene reduced in data from patients with cancer vs. healthy controls?). This enables rapid, powerful cross-characterization of the sources of variation within the dataset.

Next, the researcher might color the scatterplot by gene expression values to identify cell subpopulations based on “marker genes.” When marker genes are not known *a priori*, the researcher might then use cellxgene’s interactive computation capabilities to calculate differential expression (e.g., t-tests) between any arbitrary set of cells (e.g., “how do donors differ?” or “how is cluster 1 different from another subset of the data?”).

This process of data partitioning, cross-characterization, and inference is often repeated many times over different facets of the data. Along the way, insights about cells (observations) can be annotated within the application to export and share with collaborators and other researchers.

**Figure 1:**
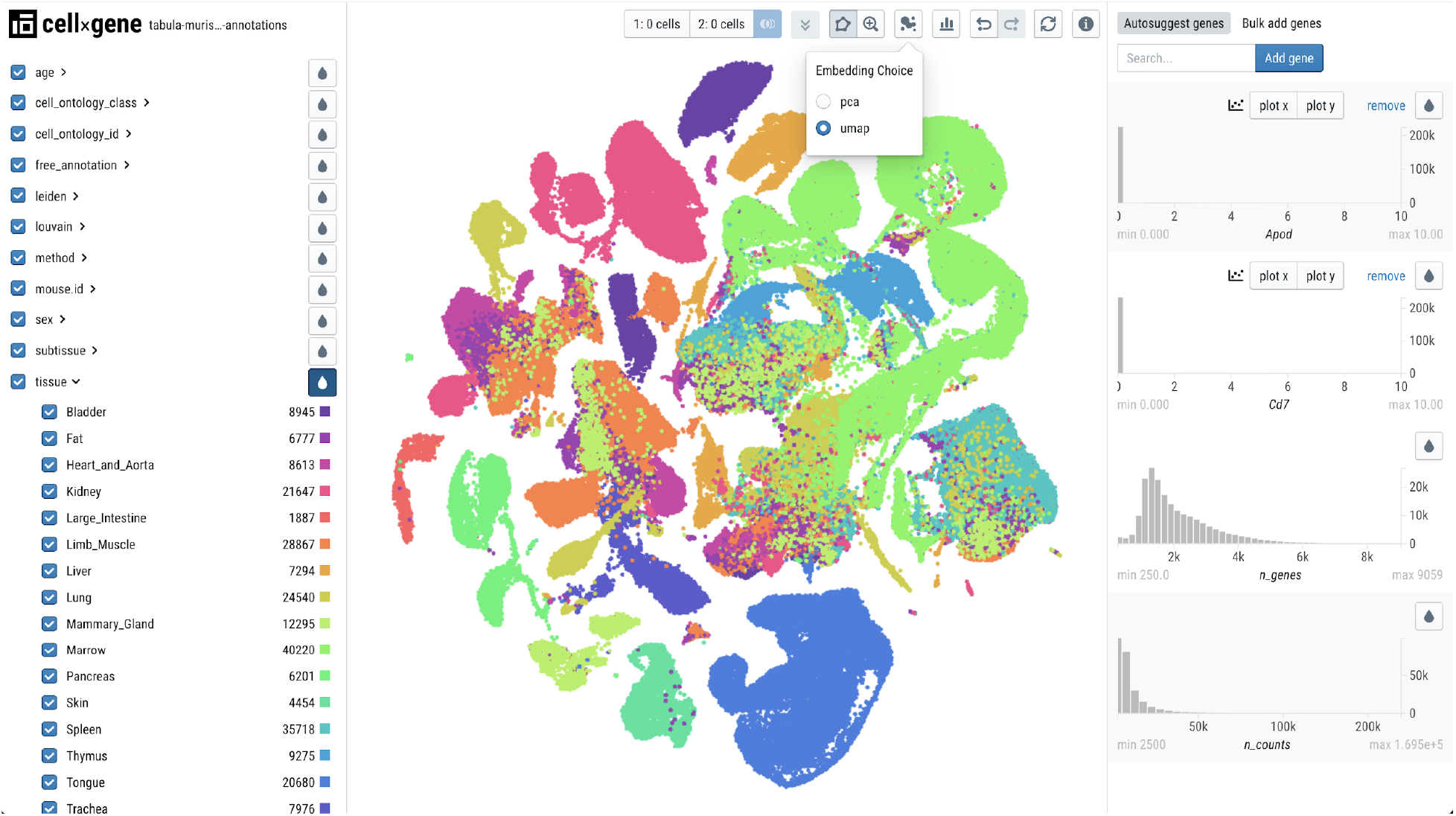
cellxgene featuring *Tabula Muris Senis* data. cellxgene is frequently used to host published and unpublished datasets for public exploration. Here, cellxgene hosts the *Tabula Muris Senis* dataset, a longitudinal mouse cell atlas (https://tabula-muris-senis.ds.czbiohub.org) [10]. Other notable examples include the COVID-19 Cell Atlas (https://www.covid19cellatlas.org) [11] and the Heart Cell Atlas (https://www.heartcellatlas.org) [9].

## Implementation and architecture

The design and technology choices in cellxgene are entirely driven by a combination of (1) requirements derived from a constant dialogue with researchers using the tool and (2) prioritizing infrastructure robustness, scalability, and speed.

To ensure that cellxgene fits smoothly into the full scientific discovery cycle, it was built to integrate with upstream and downstream tools within the computational ecosystem for single-cell analysis (e.g., ScanPy [2], Seurat [7], and Bioconductor [8]). Under the hood, however, the application implements domain-agnostic abstractions, ensuring generalizability to other single-cell data types and domains of research where scalable data exploration is required.

cellxgene’s architecture consists of a Python server and a JavaScript client. Data ingress and egress is accomplished by lazy loading of metadata, facilitated by a REST API. Transfer of data between the client and the server is handled by Google’s FlatBuffers, a binary serialization protocol [6]. cellxgene’s flexible infrastructure has enabled deployment inside of intranet environments by various institutions for large, private collaborations, and inside of platforms like Galaxy [4].

Most affordances in the interface affect other visualizations and controls (for example, filtering by metadata emphasizes corresponding items in an embedding). As such, architectural decisions had to account for caching and reconciliation of the data state upon arbitrary subsetting of data and metadata [5]. To accomplish this with millions of observations in client side JavaScript, we used a combination of D3 [1] WebGL, a purpose-built high-performance multi-dimensional filter inspired by Mike Bostock’s original crossfilter [3], and a purpose-built Pandas dataframe-like module for in-browser use [5].”

## Future Directions

cellxgene is under active development. Active areas of design and development include improvements to in-browser computation, the addition of a full workspace for recording observations about features (genes), the addition of heatmap and dot plot views that enable direct comparison of groups of observations (e.g. clusters) and groups of features (e.g. sets of genes). We are also platformizing the software to enable easy publishing and exploration of single cell datasets.

In anticipation of “atlas” sized datasets of tens of millions of cells or more, cellxgene will continue to invest in scalability (the largest dataset at time of writing is millions of cells), both in terms of systems architecture and displaying information.

## Availability and contact

cellxgene can be found at <URL>.

We appreciate and encourage contributions to or extension of cellxgene. All source code is available under an MIT license on Github at <http://github.com/chanzuckerberg/cellxgene>.

Extensive user documentation, including installation, usage, and contact information, is available at <http://chanzuckerberg.github.io/cellxgene>.

## Acknowledgments

Josh Batson, Deep Ganguli, Fabian Theis at Theis Lab, Bruce Aranow at Aronow Lab at Cincinnati Children’s Hospital, The Dana Pe’er Lab at Sloan Kettering Institute, Benjamin Humphreys at Washington University in St. Louis, Loyal Goff, Vlad Kiselev at the Wellcome Trust Sanger Institute, Valentine Svensson at Serqet Therapeutics, and Alok Saldanha at Novartis.

